# Developing an enzyme selection tool supporting multiple hosts contexts

**DOI:** 10.1101/2021.09.09.459461

**Authors:** María Camarena, Pablo Carbonell

## Abstract

Engineering biological organisms that allow the integration of alternative metabolic pathways to natural ones is one of the goals of synthetic biology. Based on this, some of the most attractive applications in terms of synthetic organisms manufacture include the production of a wide range of pharmacologically useful metabolites produced in a sustainable and environmentally friendly way. Also, the biostable molecules green-production involves different types of therapeutic processes, e.g. prostheses and grafts stabilisation. Regarding the viability of genetically modified organisms, metabolic pathways must be first properly designed, taking into consideration the type of host organism that will be involved in metabolic production, as well as its biochemical and environmental conditions. To ensure the correct growth of these synthetic organisms, the enzyme selection must guarantee both the organism survival (and proliferation) and the optimal production of the desired metabolite. Developing enzyme selection tools is essential to enhance and make cost-effective the metabolic pathways design. This technical note presents the update of Selenzyme, the enzyme selection tool which is based on organisms taxonomic compatibility and allows appropriate enzyme selection considering its amino acid sequence. The purpose of the update is to allow multiple host input, in order to perform an affinity comparison between target organisms and each host. The affinity differences will depend on which host to be considered, allowing the user to select the optimal host for the enzyme concerned.

## 1 Introduction

Traditional production modes have been reconsidered due to the context of climate change and health concerns. Governments and research institutions are applying and studying new renewable technologies, which are constantly being analyzed to achieve the best possible service. Major industries are beginning to incorporate sustainable forms of production into their manufacturing chains, thereby promoting an economic form based on products derived from renewable biological resources, thus structuring the global bioeconomy [8].

In the biocompounds production sector such as pharmaceuticals and molecules of biological interest, these processes can be carried out using techniques based on synthetic organisms. These organisms provide a renewable and sustainable molecular source, contributing to industrial sustainability and the use of green energy sources. To that end, synthetic biology provides mechanisms for gene editing and generation of organisms producing a wide range of molecules, arousing great ecological and also bio-economic benefits [4]. Such organisms can be considered a renewable source of production. In many cases, high yields and even new compounds that had not been synthesised prior to the use of these organisms can be achieved using biofactories. For these reasons, the biomanufacturing market interest has generated its upward growth in recent years, offering an optimistic environment that promotes the generation of new analyses and studies in the industry.

The study and development of techniques that contribute the synthetic biology sector boosting imply an investment in the future, in order to obtain long-lasting and sustainable production systems. Many of these techniques are based on software development to cover the following features: recording and analysing data obtained through experimental *praxis*, generating new models that are able to predict the behaviour of models, relating variables taking into account the characteristics of each one, and many other functionalities whose execution depends on the context. This technical note references one of those software-based tools. Selenzyme [3], originally developed at the Manchester SYNBIOCHEM biofoundry, is a free online enzymatic selection tool that can be used for a wide range of purposes. An additional code compatible with the original software is presented here, through which new features of Selenzyme are generated.

### 1.1 Selenzyme: enzyme selection tool

Selenzyme operates as an enzyme selection tool, as the information it provides allows the user to select the enzymes of greatest interest for the desired application, including metabolic engineering, genome annotation, disease biomarkers, biosensor development, drug design and toxicity assessment. The tool operation will be briefly explained below.

Initially, the user has to enter a SMARTS sequence (or SMIRKS) representing the biochemical transformation (coding for a desired enzyme) and a single host organism to which he/she wishes to introduce a gene coding for the desired enzyme. The tool has access to databases that relate sequences and associated catalysed reactions, as well as organisms phylogenetic classifications. Using these data, the tool provides the following data of interest in enzyme selection: phylogenetic distances between the host organism (entered by the user) and the organisms from which the gene encoding the desired enzyme will be extracted; similarity between sequences and catalysed reactions; multiple alignment to detect conserved sequences; active and catalytic regions; properties such as solubility and transmembrane regions. Thus, Selenzyme generates adequate information for the proper target organisms selection, which are directly dependent on the host organism and reaction entered by the user.

Selenzyme features provide the most relevant data for the metabolic pathways design, but these functionalities are limited to the engineering of a single organism rather than a taxonomic group, which can be a limiting factor considering the application for which the use of this tool is intended.

### 1.2 Why do we need supporting multiple hosts?

As mentioned above, the tool can be used in multiple different contexts, and supporting input from a single host could be problematic. Allowing multiple host input may be of high interest for applications that require the design of different metabolic pathways for the same enzyme. In this case, the different pathways design would depend on the different hosts entered, depending directly on the user’s multiple choice. Another possible advantage of being able to support multiple hosts would be the host organism selection, on which the experiments are to be performed prior to the physical experiments beginning (i.e., this new option would make it possible to discard experiments on host organisms that may not be feasible).

Considering the advantages that can be obtained by the user entering multiple hosts, it has been thought convenient to implement a code that solves the problem of the single host. This code will be explained in detail below.

## 2 Software implementation

This technical note presents an update of the previously published Selenzyme tool, which allows results to be obtained by entering multiple hosts. A software programme has been developed, which is consistent and compatible with the initial mode of operation of Selenzyme, as it does not modify the internal code and takes into consideration the possibility of entering a single host by the user.

In detail, the generated code operates by analysing the phylogenetic groups IDs. The target organism is considered to be the organism whose enzyme reaction is potential to be selected for introduction into the host organism. The affinity between host and target is determined by their phylogenetic distances. The phylogenetic groups to be considered are: taxon of the target organism, taxon of the host organism and, if present, taxon of the common ancestor of the target and host. A graphical scheme used to determine phylogenetic groupings is shown in figure 1. A taxon is represented by a square node, and as can be seen, two possible cases have been taken into account for the calculation of distances. Working on the assumption of figure 1.a, the distances between the host and target taxa are calculated using equation (1), where: d1 is the desired distance, d2 is the distance between the host taxon and the common ancestor taxon, d3 is the distance between the target taxon and the common ancestor taxon. In addition, a node has to be subtracted so that when the target and the host share phylogenetic group the distance is null.

**Figure 1:**
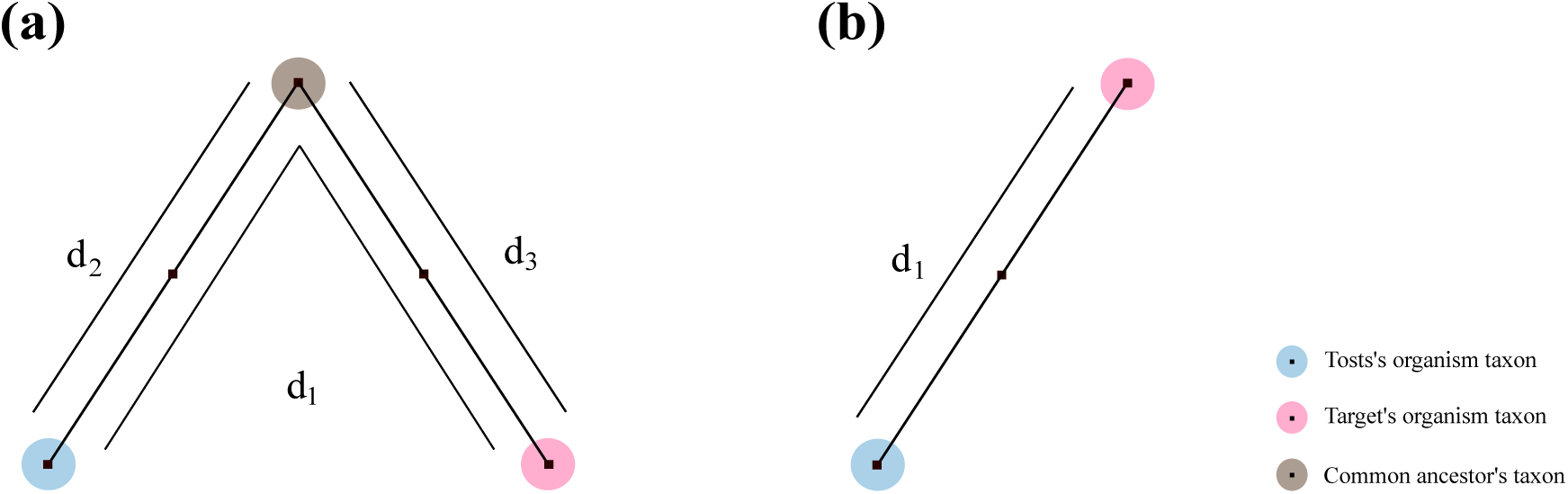
Phylogenetic distances determined by the number of taxons between host’s taxon and the taxon of the target organism. (a) Specified distances in case the host organism and the target organism share a common ancestor, (b) Specified distances if the host organism and the target organism do not share a common ancestor.

This new functionality also contemplates both negative and positive distances. Positive distances correlate distances where the host is not in the target lineage (this is the way of calculating distances originally defined in Selenzyme). An example is shown in figure 2 of a positive distance is the case of Saccharomyces cerevisiae (ID: 4932). Otherwise, negative distances correspond to those where the host and target are in the same lineage. An example of negative distance can be seen in figure 2 in the case of Bacteria (ID: 2).

**Figure 2:**
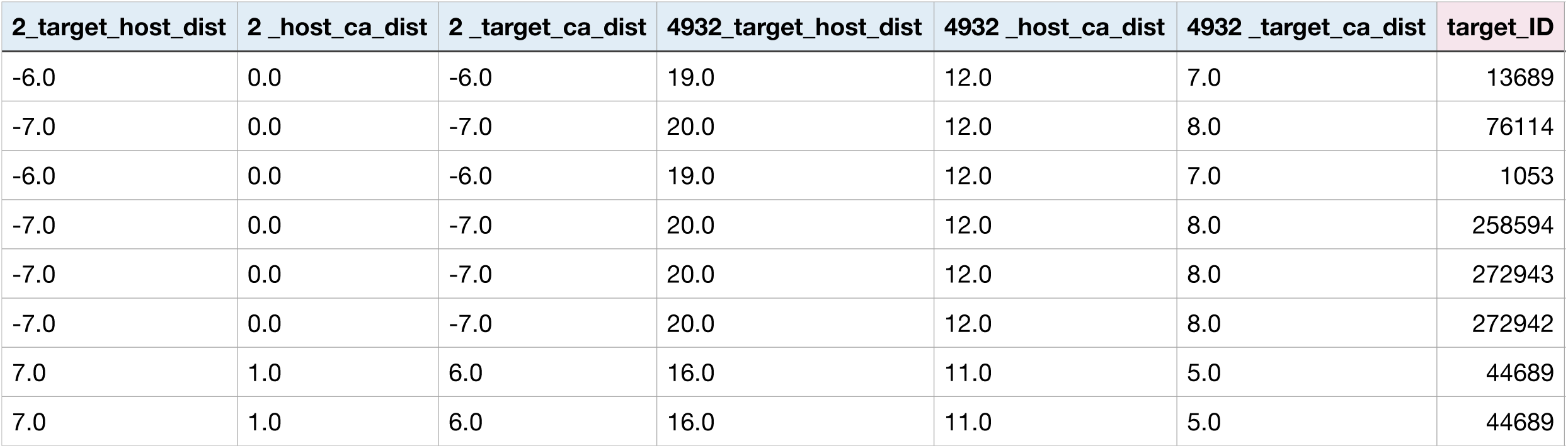
Resulting Selenzyme’s update table containing the best rated sequences for TAL enzyme. Regarding this output, both *Bacteria* (2) and *Saccharomyces cerevisiae* (4932) hosts have been entered as an array. Last column contains each target ID.

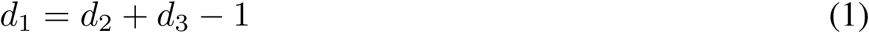

The original operation mode allows the entry of a single host, and the tool generates a csv file containing the data mentioned in the Selenzyme description section. In turn, it is possible to use a new operation mode which allows the input of multiple hosts. Figure 2 shows one example of the updated version output table. Considering the host/target common ancestor organism, the same csv file as in single host entering mode is generated, but the following columns are added for each of the introduced host organisms: phylogenetic distance between the host organism and the target organism, phylogenetic distance between the host organism and the common ancestor organism, phylogenetic distance between the target organism and the common ancestor. Each column name contains the NCBI taxonomic ID of the host, as multiple host identifiers are possible as input. In addition, another column is generated where the NCBI taxonomic ID of the target ID is provided. Notably, by using the updated code, the user can also introduce widely class host organisms, such as Gammaproteobacteria, which was not supported in the previous version of Selenzyme.

*The updating code can be found at the following URL:* https://github.com/pablocarb/selenzy/tree/newtax

## 3 Potential applications

### 3.1 Taxonomy clustering

Potential applications of the new taxonomic feature include the development of an enzyme selector based on similarity (homology) across the chemical, sequence and taxonomic spaces. Previous studies have shown that the two latter spaces can be related through evolution to the chemical space of enzyme function [2] therefore integrating the spaces might provide a more accurate selection score. Selenzyme already implemented a multiple sequence alignment (MSA), which allows clustering the sequences. Similarly, the new information about taxonomic distances for sequences can be used to cluster the sequence into those that are closer to each host. In addition, considering chemical space, Selenzyme performs the reaction similarity calculus, which allows a numerical comparison between each chemical reaction. In section 3.2 reaction similarity has been used in order to obtain reaction difference. Combining the different space information, it might be possible to select those sequences that are more relevant to each host. An example of this is explained in the section below.

### 3.2 An example using the naringenin metabolic pathway

Naringenin is a flavonoid family compound of high interest due to its pharmacological benefits, so it is currently traded with it. Therefore, and regarding design optimization, considering viable genetic circuits before plasmids assembly is considered highly relevant. Naringenin metabolic pathway consists of four enzymes, which is shown in figure 3: TAL, 4CL, CHS and CHI. Regarding to each single enzyme, the target selection can be optimized if multiple host entering is used, as the selection range covers more than one optimal possibility depending on which host is to be considered.

**Figure 3:**
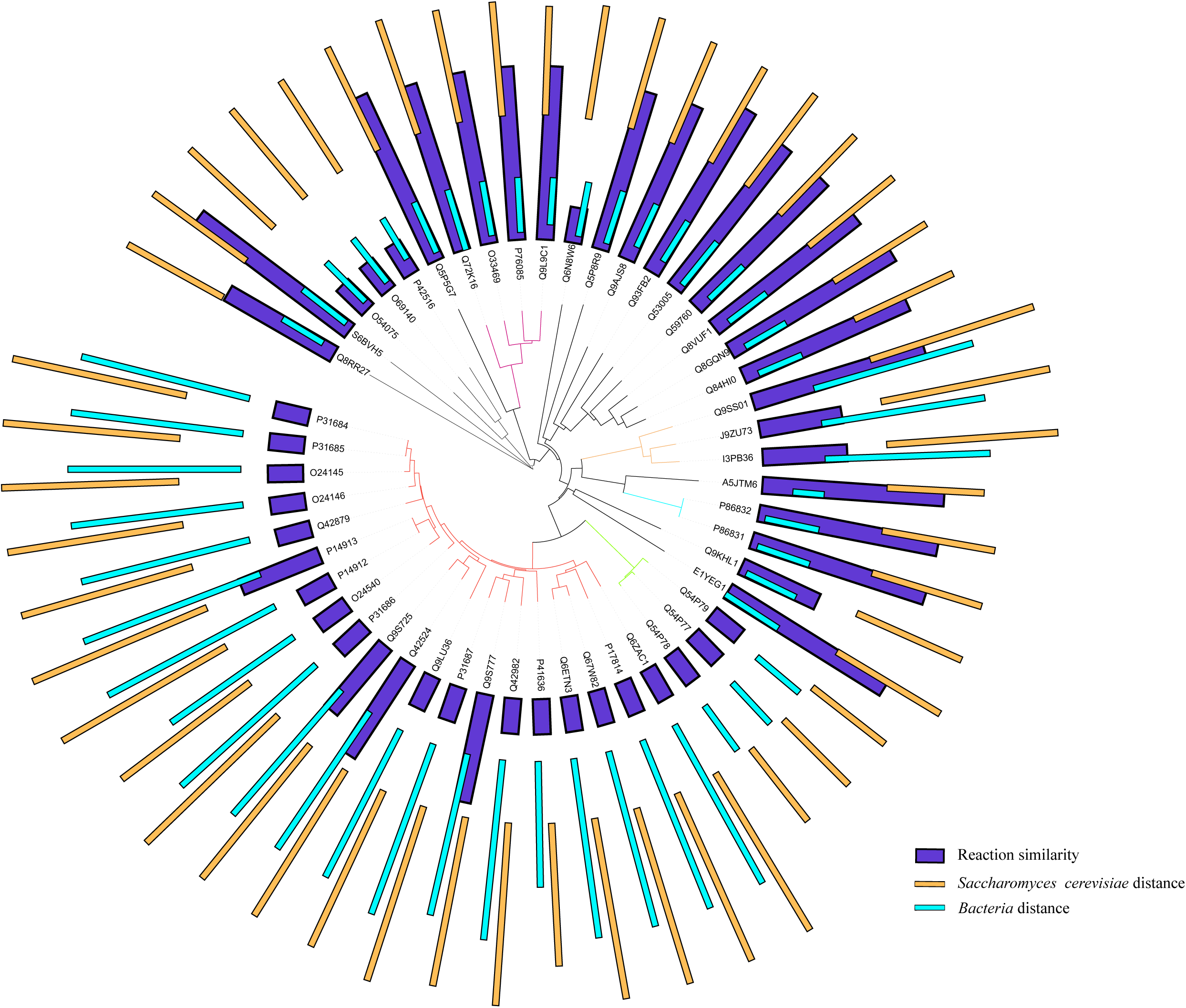
Clustered phylogenetic tree of TAL variant sequences: 50 sequences ID are represented. Each color refers to a cluster, and the black ID branches are considered as single clusters, regarding *length* criteria. In addition, simple bar chart are represented in three different colors: mulberry represent reaction similarity between the sequence and the enzyme requested; blue represent *Bacteria*’s and each target sequence ID taxonomic distance; orange represent *Saccharomyces cerevisiae*’s and each target sequence ID taxonomic distance

**Figure 4:**
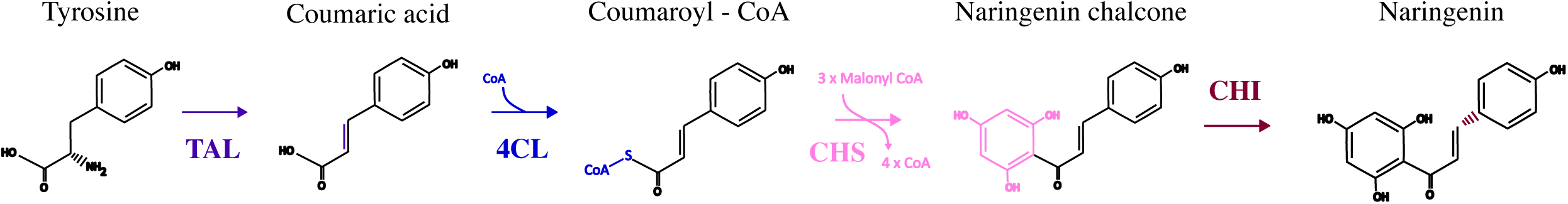
Naringenin metabolic pathway.

In this case, taxonomy clustering can be considered as a computational advantage. Moreover, we wanted to graphically represent the differences that can be found between each target regarding the chemical, sequence and taxonomic spaces mentioned in section 3.1. For each cluster, one of the ID sequences can be considered depending on its taxonomic similarity with the each host entered. Figure 3 shows the TAL sequences clustered phylogenetic tree generated using [iTOL](https://itol.embl.de/), Interactive Tool Of Life, an online tool that allows phylogenetic trees to be generated and modified. Regarding the clustering, TreeCluster tool explained in [1] was used to generate the cluster, in which *length* method with t = 0.3 has been used. Clustering criteria has been selected regarding tree leaves length. The clustering method used to acquire figure 5 is *length* method, in which “the cluster does not contain any edges above length *t*”. Each method required the entering of a threshold *t*; for this case *t = 0*.*3*.

**Figure 5:**
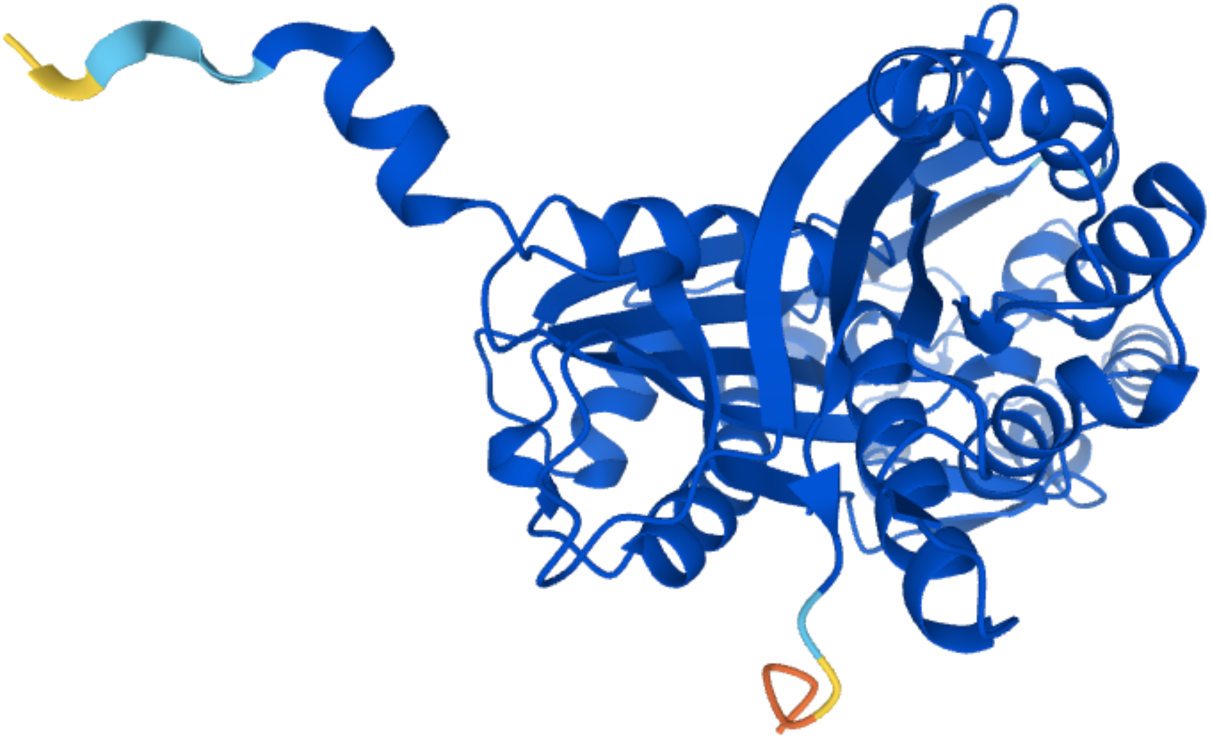
Naringenin Chalcone synthase 1 3D structure, obtained from [AlphaFold](https://alphafold.ebi.ac.uk/) database.

As mentioned above, chemical, sequence and taxonomic spaces of each enzyme have to be considered in order to correctly characterise the enzyme choice. To this end, and taking into account each cluster, it was expected that the properties of each of the spaces should be similar in the same cluster. However, when graphically representing the data, some differences were found. Clustering represents the sequence space. The bar charts shown in figure 5 will be analysed below.

Regarding chemical space, purple bar charts are represented. Each bar is assigned to a target, and the length of each bar represents the reaction difference between requested and target enzymes. Reaction difference (*R*_*d*_) is determined by using equation 2:

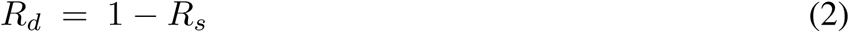

where *R*_*s*_ is reaction similarity between the reactions catalysed by target and requested enzymes. *R*_*s*_ parameter is obtained from Selenzyme’s table output. As shown in Figure 5, there are differences in four of the twenty chemical reactions within the red cluster. Similarly, orange cluster presents differences in one of three represented by its cluster. The remaining clusters show homogeneous parameters within the same cluster.

Considering taxonomic distances between hosts and targets from each cluster, taxonomic space can be characterised. Blue bars represent *Bacteria*’s distance between each target organism (from where the sequence is provided), and as shown it can be positive or negative distance depending on the taxonomic relative positions. Selenzyme’s update allow negative distances, as explained previously, as most of the target share *Bacteria*’s lineage. Orange bars, in contrast, represent *Saccharomyces cerevisiae*’s distances. As shown, a certain homogeneity can be considered within each cluster. Taxonomic spacing is relevant, as enzyme production will be higher the closer two organisms are to each other.

### 3.3 Future directions

Each enzyme owns its 3D structure, which also determines both functionality and work-way. Therefore, considering the structural space is a relevant point in order to enhance Selenzyme’s features. A new structural score can be implemented considering the protein structure provided by AlphaFold, the protein structure database developed by DeepMind and the EBI. This database obtains 3D protein structures by using AI and machine learning algorithms that work highly accurately. The structure score could be implemented considering advanced studies, such as molecular dynamics, docking, hotspot identification or even smart library designs. Hence, the functional score would provide key features for the final enzyme selection, as it would represent physical and functional enzyme performance (that also depends on its interaction with the host medium).

Considering one type of enzyme, studying the trade-off between its sequence expression and its dynamic response is a key factor to accurately tune the genetic circuit performance.

Expression efficiency, on which depends the enzyme concentration, is related to the phylogenetic distance between the host organism and the target one [5]. More phylogenetic distance between the organisms can imply lowest expression values of the enzyme, as the differences in each organism medium increase with the distance and hampers the expression viability. On another note, with regard to the dynamic response, enzymatic efficiency depends on its kinetic parameters, such as km and kcat. Considering one type of enzyme, different enzyme variants have different kinetic parameters. Consequently, and for a given study, specific kinetic parameters will be of relevance and these will be determined by a given enzyme variant. Considering both scenarios, this type of studies can bring substantial enhancements in enzymatic selection.

To implement novel kinetic studies, enzymatic kinetic parameters need to be used, and these can be obtained from [BRENDA’s](https://www.brenda-enzymes.org/) database [6]. Unfortunately, kinetic parameter data are scarce and not available for any given enzyme. To cover the lack of data and therefore allow novel kinetic studies implementation, machine learning algorithms can be developed in order to predict kinetic parameters values [7].

### 3.4 Conclusions

In summary, in order to improve the selection of enzyme sequences for metabolic pathway design, we have introduced here a new feature that allows the enzymatic selection based on multiple host contexts. Combining this new feature with other future functionalities like AI-based prediction, will allow a more precise selection of functional sequences potentially leading to an improved pathway performance.

## 3.5 Acknowledgments

MC was supported by an AI2-UPV studentship. PC acknowledges funding from UPV Talento Programme and European Union’s Horizon 2020 research and innovation programme under grant agreement number 814408 (ShikiFactory100). This research was partially supported by grants MINECO/AEI, EU DPI2017-82896-C2-1-R and MICINN/AEI, EU PID2020-117271RB-C21.

